# Geraniol Enhances Inhibitory Inputs to Paraventricular Thalamic Nucleus and Induces Sedation in Mice

**DOI:** 10.1101/2021.04.25.441289

**Authors:** Ling Xu, Yan Wang, Ya-Yue Yang, Xiao-Xiao Hua, Li-Xia Du, Jian-Yu Zhu, Li-Na Huang, Fang Fang, Ming-Zhe Liu, Rui Zhang, Jin-Bao Li, Yan-Qing Wang, Ling Zhang, Wen-Li Mi, Di Mu

**Affiliations:** Department of Anesthesiology, Shanghai General Hospital, Shanghai Jiao Tong University School of Medicine, Shanghai 201620, China; Department of Integrative Medicine and Neurobiology, School of Basic Medical Science; Institutes of Integrative Medicine; Institutes of Brain Science, Shanghai Medical College, Fudan University, Shanghai 200032, China; The First Rehabilitation Hospital of Shanghai, Tongji University School of Medicine, Shanghai 200090, China; Department of Endocrinology, Shanghai General Hospital, Shanghai Jiao Tong University School of Medicine, Shanghai 201620, China; Department of Respiratory, The First Affiliated Hospital of Guangzhou Medical University, Guangzhou 510120, China

**Keywords:** Geraniol, Sedation, paraventricular thalamic nucleus, tonic inhibition, GABA_A_ receptors

## Abstract

Geraniol (GE), a plant-derived acyclic monoterpene, shows a wide variety of beneficial effects. Notably, recent studies have reported the potential sedative effects of GE in fish and rats. However, the mechanisms of GE in sedation remain elusive. Here, we found that GE reduced locomotion, relieved pentylenetetrazol (PTZ)-induced seizures, altered the electroencephalogram (EEG), and facilitated general anesthesia in mice. Meanwhile, GE decreased c-Fos expression and suppressed the calcium activity in the paraventricular thalamic nucleus (PVT). Microinjection of GE into the PVT reduced locomotion and facilitated propofol-induced anesthesia. Furthermore, the electrophysiology results showed that GE-induced dramatic membrane hyperpolarization and suppressed the neuronal activity of PVT neurons, mainly by prolonging spontaneous inhibitory postsynaptic currents and inducing tonic inhibitory currents via GABA_A_ receptors. Our study revealed that GE enhances inhibitory inputs to PVT neurons and induces sedation in mice. These findings provide a potential candidate for further development of sedatives and anesthetics.

**Graphic Summary:** 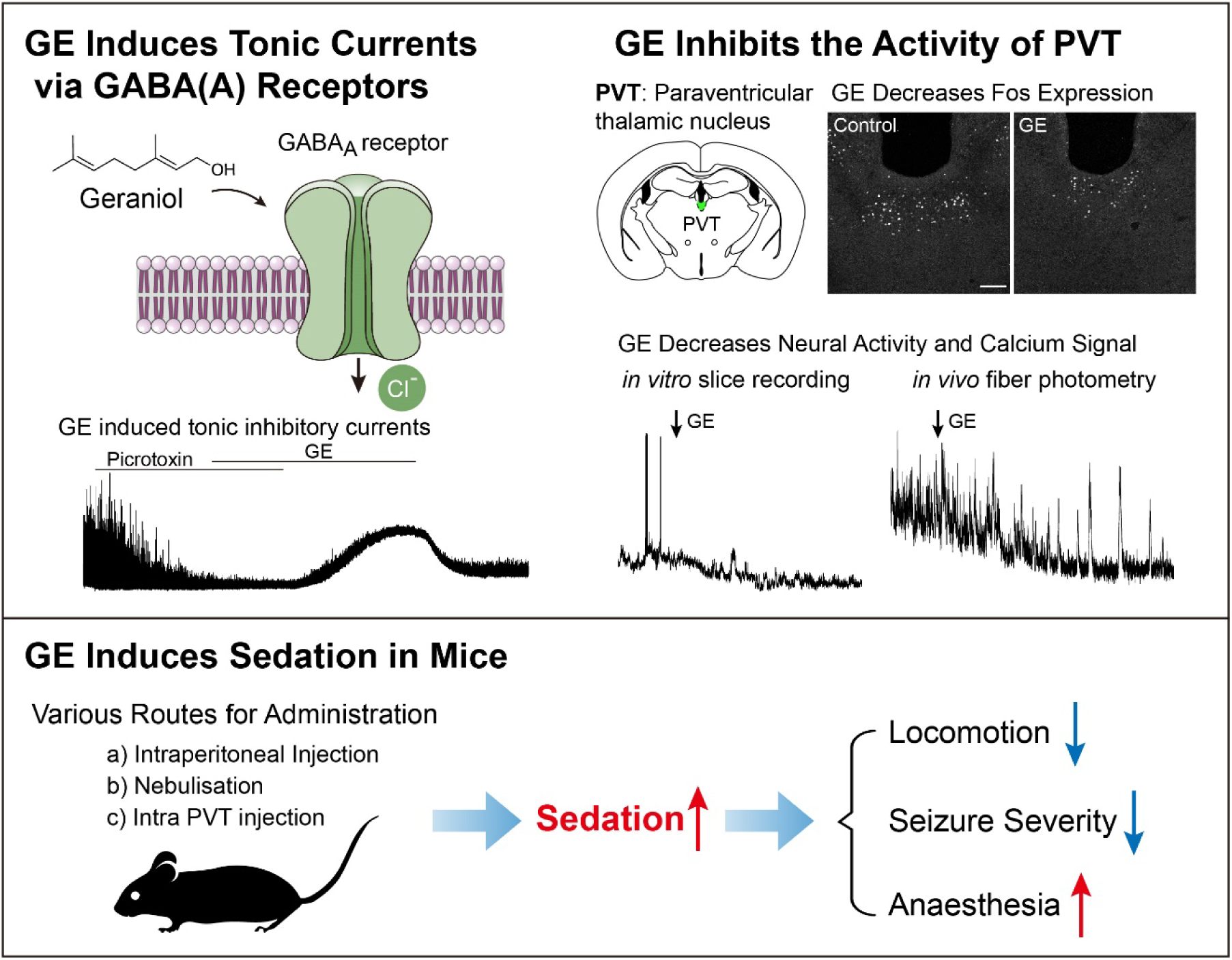

## Introduction

Sedatives are depressants that reduce the irritability or excitement of the central nervous system. They are mainly used in surgical operations or intensive care units with analgesics and muscle relaxants to achieve the “triad of anesthesia” (Reade and Finfer, 2014). Sedation with propofol without intubation is widely used in both first and second-trimester surgical abortions in the outpatient setting (Dean et al., 2011). Moreover, there is an increasing need for sedation in pediatric patients (Coté and Wilson, 2019). Dissecting the mechanisms of sedation and developing novel sedatives have been the focus of sedation and anesthesia.

Plant extracts with sedative effects have long clinical experience in insomnia and epilepsy (Gaston and Szaflarski, 2018; Shi et al., 2016). Geraniol (GE; 3,7-dimethylocta-trans-2,6-dien-1-ol), an acyclic monoterpene, is abundant in essential oils extracted from lemongrass, rose, lavender, and other aromatic plants (Lapczynski et al., 2008; Pavan et al., 2018). It is classed as a safe flavor ingredient by the FDA and has an IFRA (International Fragrance Association) standard (Lapczynski et al., 2008). Reports showed that GE had anti-microbial, anti-inflammatory, anti-oxidant, anti-nociceptive, neuroprotective, and anti-cancer effects (Cho et al., 2016; Khan et al., 2013; La Rocca et al., 2017; Lv et al., 2017; Rekha et al., 2013; Thapa et al., 2012). Notably, it might also have sedative effects in fish and rats (Can et al., 2019; Medeiros et al., 2018). However, the underlying mechanisms of GE in sedation remain elusive, and the role of GE in these processes requires further dissection.

Sedatives have robust locomotor sedating effects (McOmish et al., 2012; Ralvenius et al., 2016) and anti-convulsant effects (Brohan and Goudra, 2017). A previous study found that GE could reduce locomotion and increased barbiturate-induced sleeping time in rats, reflecting the depressant effect of GE (Medeiros et al., 2018). Pentylenetetrazole (PTZ) is a central nervous system convulsant being thought to inhibit GABA_A_-mediated Cl^−^ currents (Huang et al., 2001). A single injection of PTZ is a commonly used acute seizure model in mice (Li et al., 2012; Van Erum et al., 2019). Whether GE could relieve the acute seizure in mice should be determined.

The corresponding brain regions and receptors involved in GE are currently unknown. A recent study found that the paraventricular thalamic nucleus (PVT) is a critical node for controlling wakefulness in mice (Ren et al., 2018). Activation of PVT neurons induces the transition from sleep to wakefulness. Conversely, suppression of the PVT neurons causes a reduction in wakefulness (Ren et al., 2018). Furthermore, previous studies have shown that GE might act on voltage-gated potassium channels (Ye et al., 2019) or calcium channels (El-Bassossy et al., 2016). A computational study revealed that the GABA_A_ receptor α1 and β1 subunits might also be putative targets of GE (Zhang et al., 2019).

In the current study, we tested the hypothesis that GE plays a vital role in inducing sedation by multiple behavioral tests and EEG recordings. Next, we determined the mechanism of GE on PVT neurons by using calcium imaging, pharmacology, and brain slices electrophysiology. Our study revealed the sedative effects of GE via acting on the GABA_A_ receptor in the PVT, providing a basis for further investigating essential oils’ mechanisms and developing novel sedatives.

## Results

### Geraniol reduces locomotion and relieves PTZ-induced seizures in mice

Inhalation is an ideal administration route according to the bioavailability and therapeutic efficacy. First, we examine whether the nebulization of GE could suppress locomotion. We kept the mice in the induction chamber and nebulized GE (1.5% in ddH_2_O, 65 ml) for 40 minutes and then conducted the open field test (OFT, *Figure 1A*). The total distance and the move duration of GE-treated mice were significantly reduced compared with the ddH_2_O-treated mice (*Figure 1B-D*), while the velocity was unaffected (*Figure 1E*). Next, we intraperitoneally injected different doses of GE (100, 200, 400 mg/kg in corn oil) and performed the OFT (*Figure 1F*). Similarly, GE reduced the total distance and move duration dose-dependently, without affecting the velocity (*Figure 1G-I, Figure 1-video 1*). These data confirm that GE has locomotor sedating effects in mice.

**Figure 1.**
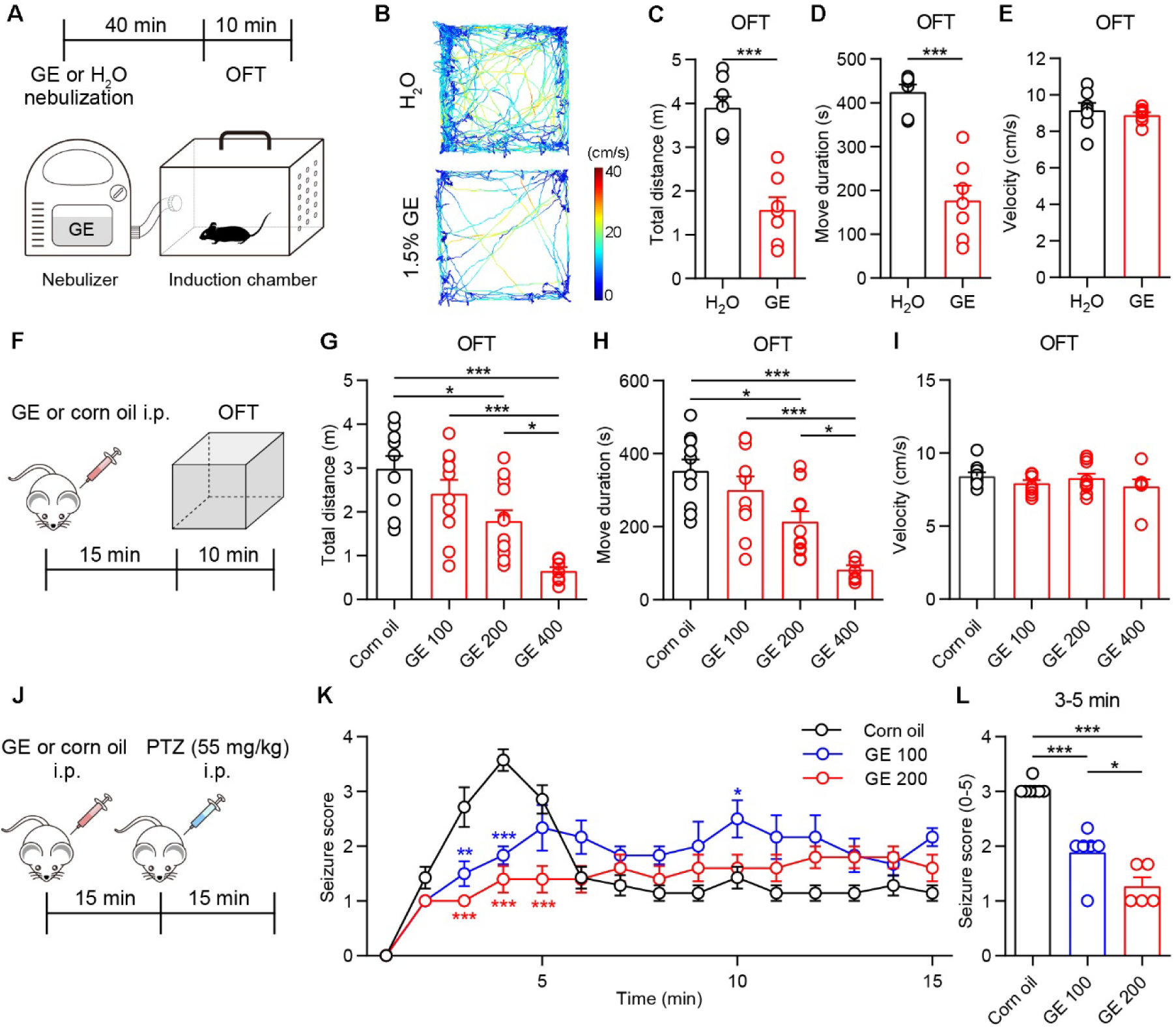
Geraniol reduces locomotion and relieves PTZ-induced seizures in mice. (A) Timeline for geraniol (GE) nebulization and open field test (OFT). (B) Representative moving tracks from an H_2_O nebulization mouse and a GE nebulization mouse. (C-E) The total distance (C), move duration (D), and velocity in the open field test, *n* = 6 mice. (F) Timeline for GE intraperitoneal injection and open field test. (G-I) The total distance (G), move duration (H), and velocity (I) in the open field test at doses of 100, 200, 400 mg/kg GE and corn oil, *n* = 7-12 mice. (J) Timeline for GE injection and PTZ (55 mg/kg)-induced seizures experiment. (K) Time course of mean scores of seizures induced by intraperitoneal injection of PTZ with GE or corn oil. Seizures symptoms were scored every 1 min for 15 min, *n* = 5-7 mice. (L) Quantification of mean score values for 3-5 min in each group. *p < 0.05, **p < 0.01, ***p < 0.001. All data were represented as mean ± SEM. Unpaired *t*-test for C, D. One-way ANOVA with Bonferroni post hoc test for G, H, K, and L. The following figure supplements are available for figure 1: **Video 1.** Intraperitoneal injection of GE and open field test. Mice were intraperitoneally injected with corn oil or GE (100, 200, 400 mg/kg). Fifteen minutes later, mice were tested in the OFT for 10 minutes. The video was played by 4x speed. **Video 2.** Intraperitoneal injection of GE and PTZ-induced seizure test. Mice were intraperitoneally injected with corn oil or GE (200 mg/kg). Fifteen minutes later, mice were intraperitoneally injected with PTZ (55 mg/kg) and videotaped for 15 minutes. The video showed the first 5 minutes after PTZ injection.

To further examine GE’s anti-convulsant effect, we used pentylenetetrazol (PTZ)-induced acute seizures. Three groups of mice were intraperitoneally injected with corn oil, 100 mg/kg GE, or 200 mg/kg GE, respectively, and 55 mg/kg PTZ 15 minutes later (*Figure 1J*). Behaviors were videotaped and manually scored from 0 (no abnormal behavior) to 5 (death) (Li et al., 2012; Van Erum et al., 2019). We found that the PTZ-induced seizure severity in the first 5 minutes was dramatically decreased in GE-treated mice compared with corn oil-treated mice (*Figure 1K, Figure 1-video 2*). Moreover, we analyzed the average seizure score of 3-5 minutes after PTZ injection, the period with the most severe symptoms. We found that the seizure score in the 100 mg/kg GE group (1.9 ± 0.2) was about two-thirds of that in the corn oil group (3.0 ± 0.1), while one-third for the 200 mg/kg GE group (1.3 ± 0.2) (*Figure 1L*). These results demonstrate that GE relieves the PTZ-induced seizures in mice, indicating the anti-convulsant effect of GE.

Next, we performed EEG recording and analyzed the power spectral density (PSD) for different frequency bandwidths. We recorded electroencephalogram (EEG) signals for 20 minutes for baseline and then intraperitoneally injected GE (200 mg/kg) or corn oil. Ten minutes later, EEG was recorded for another 20 minutes. We found that GE significantly enhanced the PSD of delta waves (0.4-4 Hz) and theta waves (4-7 Hz) (*Figure 2A-E*), while corn oil showed no apparent effect for all bandwidths (*Figure 2F-J*). These data indicate that GE enhances both delta and theta waves in mice.

**Figure 2.**
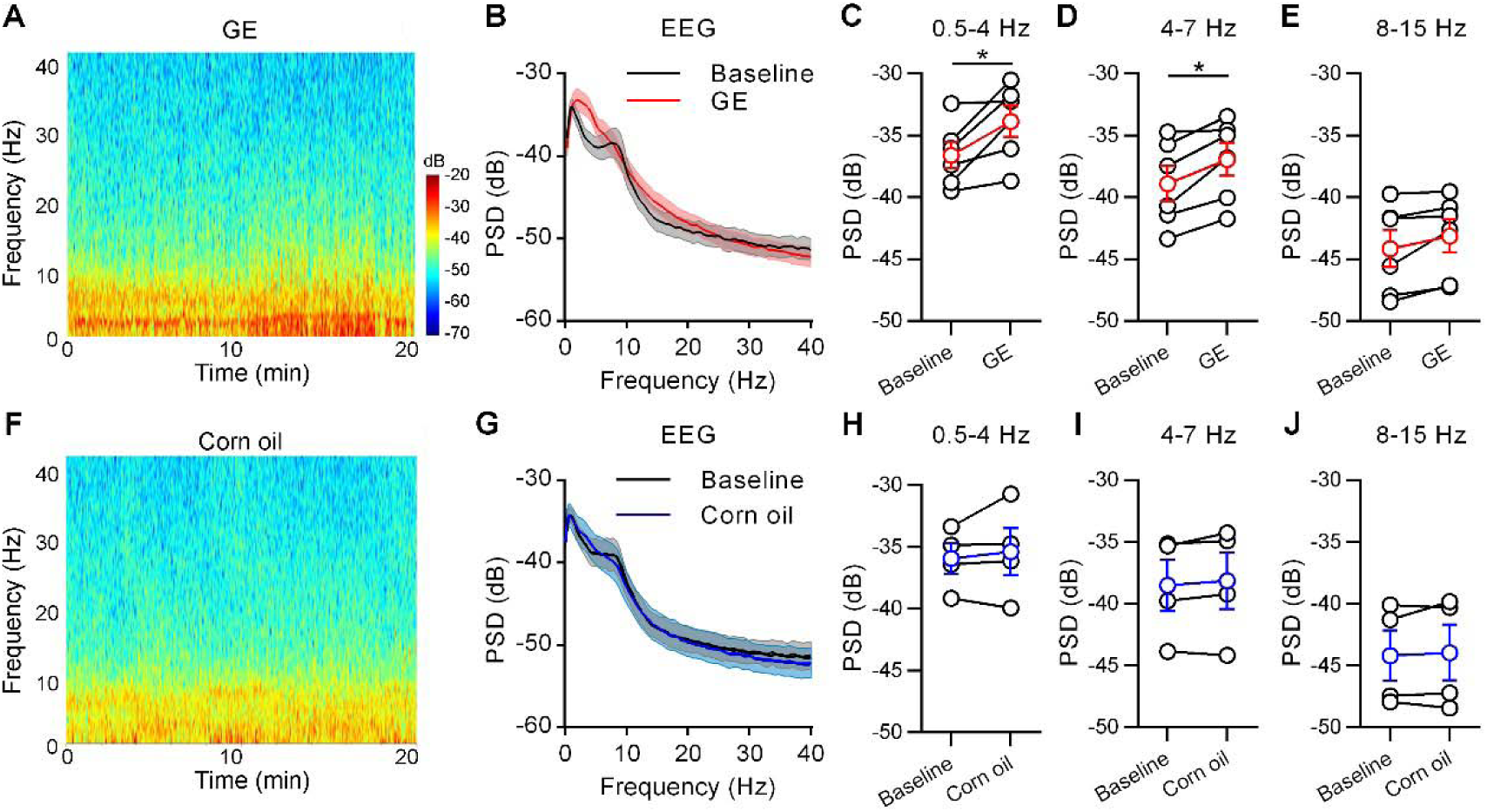
Geraniol alters the EEG in mice. (A) Representative power spectral density (PSD) of EEG data of a GE-injected mouse. Warm colors (red) represent higher power, while cool colors (blue) represent lower power. (B) Power spectra before and after GE injection, *n* = 6 mice. (C-E) Quantification of the PSD for delta waves (0.5-4 Hz) (C), theta waves (4-7 Hz) (D), and alpha waves (8-15 Hz) (E). (F) Representative power spectral density of EEG data of a corn oil-injected mouse. (G) Power spectra before and after corn oil injection, *n* = 4 mice. (H-J) Quantification of the PSD for delta (0.5-4 Hz) (H), theta (4-7 Hz) (I), and alpha (8–15Hz) (J). *p < 0.05. All data were represented as mean ± SEM. Paired *t*-test for C and D.

### Geraniol facilitates general anesthesia

We next examine whether GE could induce anesthesia. We nebulized GE or intraperitoneally injected GE (200 mg/kg or 400 mg/kg) and found that GE could not directly induce the loss of righting reflex in mice (data not shown). And then, we explore the potential role of GE in facilitating anesthesia. We first nebulized GE for 40 minutes and then intravenously injected propofol (*Figure 3A*), and we recorded the time of loss of the righting reflex (LORR) and return of the righting reflex (RORR), which have been used as a surrogate measure for the loss and resumption of consciousness under anesthesia (Franks, 2008). We found that propofol (PRO, 20 mg/kg) led to a 100% LORR rate within 5 s in both GE and control groups. Nebulization of GE increased the propofol-induced RORR compared to the ddH_2_O (*Figure 3B*). Consistently, intraperitoneal injection of GE dose-dependently increased the propofol-induced RORR compared to corn oil (*Figure 3C-D*). To further determine whether GE could reduce propofol dosage to achieve the same RORR, we reduced the dose of propofol from 20 mg/kg to 15 mg/kg. The RORR in the 200 mg/kg GE + 15 mg/kg PRO combination group was comparable to that in the corn oil + 20 mg/kg PRO group (*Figure 3E*), indicating that GE could reduce the dose of propofol for safe anesthesia.

**Figure 3.**
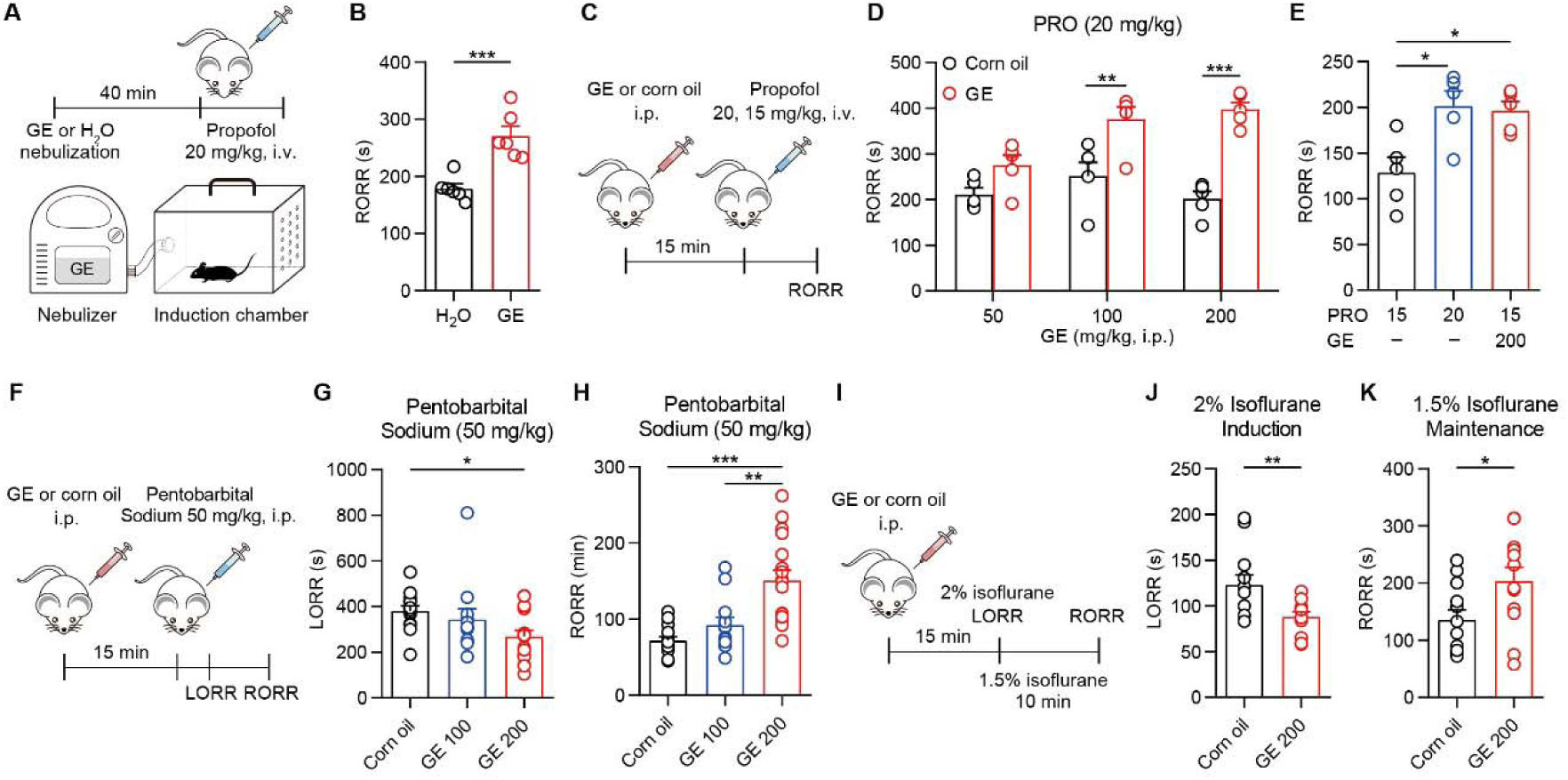
Geraniol facilitates general anesthesia. (A) Timeline for GE nebulization and open field test. (B) RORR in propofol (20 mg/kg, i.v.)-induced anaesthesia with GE nebulization (1.5% in ddH_2_O, 65 ml in 40 minutes), *n* = 6 mice. (C) Timeline for GE intraperitoneal injection and propofol-induced anesthesia experiment. (D) RORR in propofol (20 mg/kg, i.v.)-induced anaesthesia with GE (50, 100, 200 mg/kg, i.p.), *n* = 5 mice. (E) RORR in 15 mg/kg propofol-induced anesthesia, 20 mg/kg propofol-induced anesthesia, and 15 mg/kg propofol combined with 200 mg/kg GE-induced anesthesia. *n* = 5 mice. (F) Timeline for GE intraperitoneal injection and pentobarbital sodium (50 mg/kg, i.p.)-induced anesthesia experiment. (G and H) LORR (G) and RORR (H) in pentobarbital sodium-induced anesthesia with GE (200 mg/kg, i.p.), *n* =12-17 mice. (I) Timeline for GE intraperitoneal injection and isoflurane-induced anesthesia experiment. (J and K) LORR (J) and RORR (K) in 2% isoflurane-induced anaesthesia with GE (200 mg/kg, i.p.), *n* = 12 mice. *p < 0.05, **p < 0.01, ***p < 0.001. All data were presented as mean ± SEM. Unpaired *t*-test for B, J, K. Two-way ANOVA with Bonferroni post hoc test for D. One-way ANOVA with Bonferroni post hoc test for E, G, H. The following figure supplement is available for figure 3: **Figure supplement 1.** GE does not impair learning and memory in propofol-induced anesthesia of mice.

We then examined the effect of GE in the general anesthesia induced by intraperitoneal injection of pentobarbital sodium or inhalation of isoflurane (*Figure 3F-K*). The results showed that GE (200 mg/kg, i.p.) reduced the LORR and prolonged the RORR in pentobarbital sodium-induced anesthesia (*Figure 3F-H*). GE also decreased the LORR in 2% isoflurane induction and prolonged the RORR after 10 minutes of 1.5% isoflurane maintenance (*Figure 3I-K*). These data indicate that GE facilitates both intraperitoneal and inhalation anesthesia in mice.

One critical concern for the side effects of anesthetics is respiratory suppression. We examined the effects of GE (200 mg/kg, i.p.) on respiratory rate and peripheral capillary oxygen saturation (SpO_2_) when combined with propofol (20 mg/kg, i.v.)-induced anesthesia. We found that GE/propofol combination did not affect the respiratory rate and SpO_2_ (*Figure 3–figure supplement 1A-B*), indicating that GE facilitates propofol anesthesia without respiratory suppression. Another concern of anesthesia is the potential impacts on learning and memory. We then adopted the Morris water maze (MWM) and fear conditioning (FC) paradigms. Three groups of mice were injected with corn oil (i.p., corn oil), propofol (20 mg/kg, i.v., PRO), or GE with propofol (GE 200 mg/kg, i.p., propofol 20 mg/kg, i.v., PRO+GE), respectively. Twenty-four hours later, the mice underwent training to find the platform for five successive days (Day 1-5, *Figure 3–figure supplement 1C*). On day 6, the platform was removed, and all mice took the probe test (*Figure 3–figure supplement 1D*). We found that all groups showed similar spatial learning efficiency and memory performance. In the FC paradigm, the mice underwent FC training 24 hours after drug injection, and they showed similar freezing times in the five conditioning trials (*Figure 3–figure supplement 1E*) and the following cue test (*Figure 3–figure supplement 1F*). These results illustrate that GE does not impair learning and memory in propofol-induced anesthesia of mice.

### PVT is a potential brain nucleus for GE’s sedative effects

To dissect the brain nucleus affected by GE, we first performed the c-Fos staining in the brain. The PVT is a critical wakefulness-controlling nucleus in the thalamus (Ren et al., 2018). We found that the c-Fos expression in the PVT was significantly decreased at 2 hours after GE treatment (200 mg/kg, i.p., *Figure 4A-C*), indicating that GE suppressed the PVT activity. We further performed *in vivo* fiber photometry to record the calcium signal of the PVT neurons. We injected the AAV-hSyn-GCaMP6s virus and implanted the optic fiber in the PVT (*Figure 4D-E*). The results showed that the photobleaching was not significant during the 20 minutes recording in the corn oil group. However, GE significantly decreased the calcium signal of the PVT (*Figure 4F-H*). These staining and *in vivo* calcium imaging results indicate that GE suppresses the activity of PVT.

**Figure 4.**
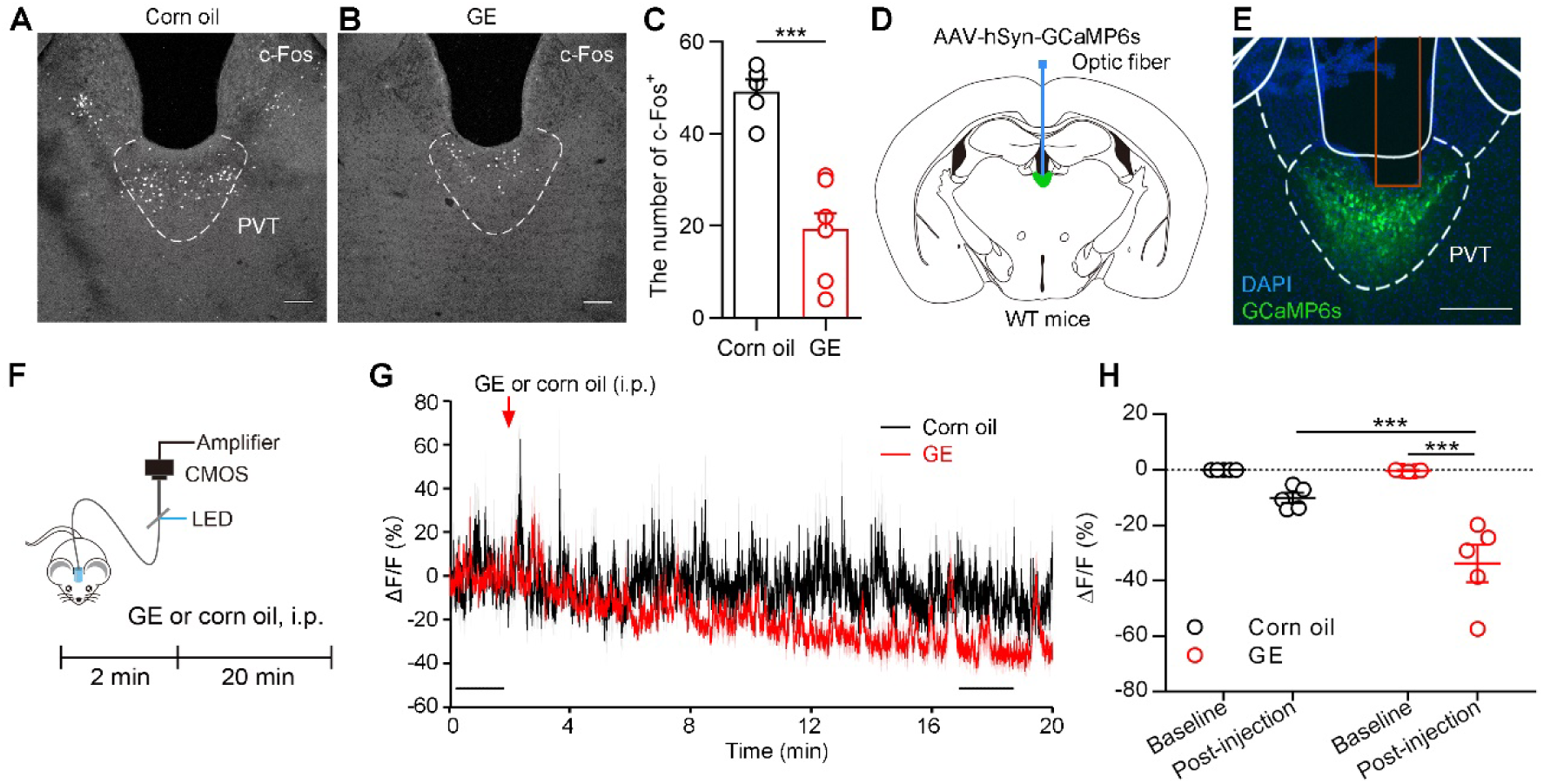
Geraniol suppresses the activity of the PVT. (A and B) The c-Fos expression in the PVT after corn oil (A) or GE injection (B). Scale bar: 100 μm. (C) Quantification of the c-Fos^+^ neurons in the PVT, *n* = 5-6 mice. (D) The schematic for virus injection of AAV-hSyn-GCaMP6s into the PVT and placement of the optic fiber above the PVT. (E) The AAV-hSyn-GCaMP6s virus expression and the fiber track (red rectangle) in the PVT. Scale bar: 200 μm. (F) Timeline for GE or corn oil intraperitoneal injection and calcium recording. (G) Averaged PVT calcium activities of GE-injected mice (red) and corn oil-injected mice (black). Two black bars represented the 2-minute baseline period and the 2-minute post-injection period (15 minutes after GE injection), *n* = 5 mice. (H) Quantification of the fluorescence intensities in the baseline and post-injection period. ***p < 0.001. All data were presented as mean ± SEM. Unpaired *t*-test for C. Two-way ANOVA with Bonferroni post hoc test for H.

### Microinjection of GE in PVT reduces locomotion and facilitates anesthesia

To further investigate the role of PVT in GE-induced sedation, we implanted cannulas in the PVT and microinjected GE or artificial cerebrospinal fluid (ACSF) into the PVT (*Figure 5A*). First, we examined propofol-induced anesthesia after microinjection of GE (1 mM, 200 nl, *Figure 5B*, top) and found that GE markedly prolonged the RORR (*Figure 5C*). Next, we used the open field test to examine the effect of PVT microinjection of GE in locomotion (*Figure 5B*, bottom). Consistent with the systemic administration, the results showed that total distance and move duration were decreased in the GE group compared to the ACSF group (*Figure 5D-F*), with the velocity unaffected (*Figure 5G*). These data demonstrated that microinjection of GE into the PVT facilitates propofol anesthesia and reduces locomotion.

**Figure 5.**
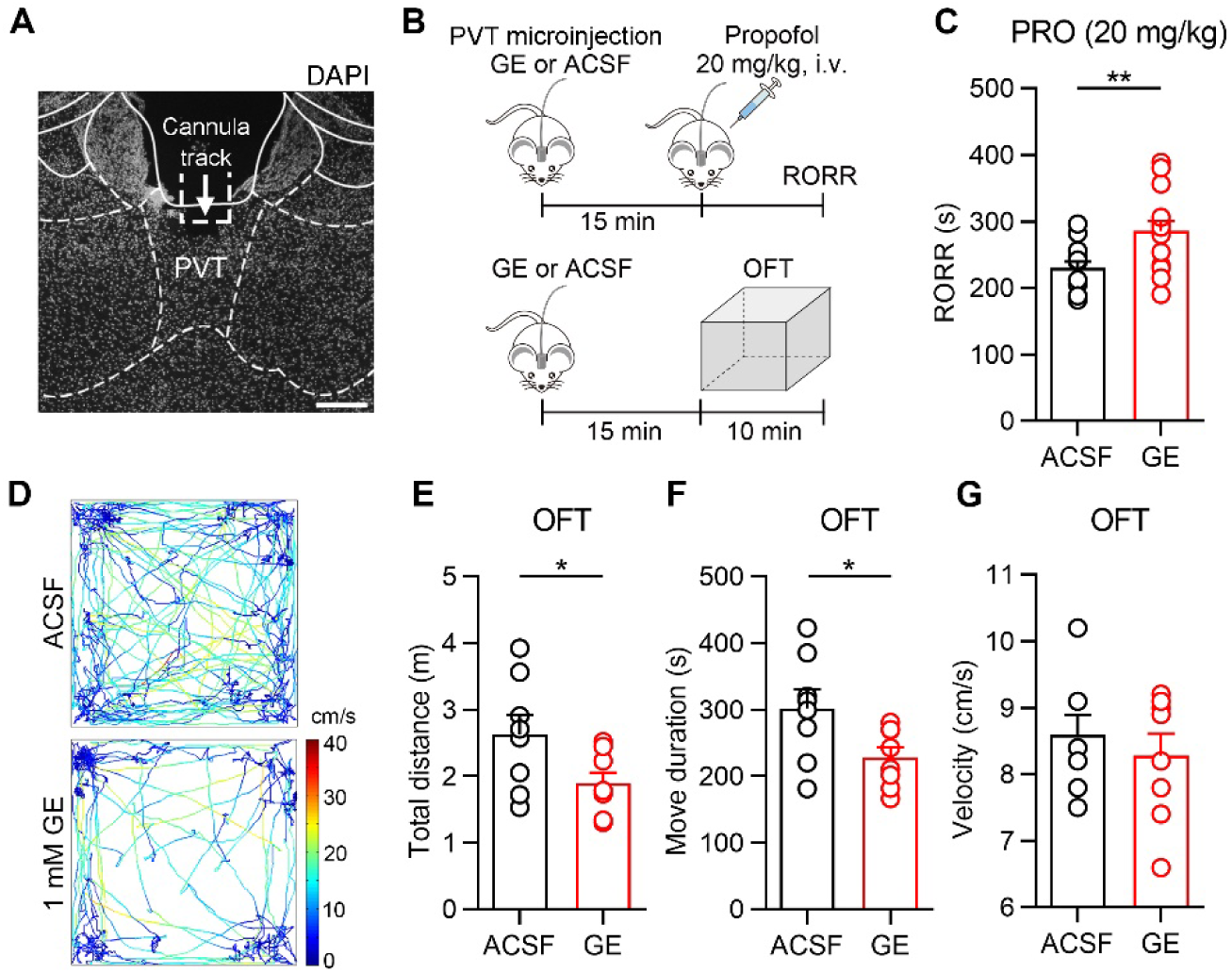
Microinjection of GE in PVT facilitates anesthesia and reduces locomotion. (A) Representative DAPI staining image from one mouse implanted with the cannula in the PVT. The arrow and dotted rectangle indicate the cannula track. Scale bar, 200 μm. (B) Timelines for GE (1 mM, 200 nl) or ACSF microinjection in propofol-induced anesthesia experiment (top) and open field test (bottom). (C) Propofol-induced RORR in GE or ACSF microinjection experiments, *n* = 14-15 mice. (D) Representative moving tracks from an ACSF (top) and a GE microinjected mouse (bottom). (E-G) Total distance (E), move duration (F), and velocity (G) in the open field test, *n* = 5 mice. *p < 0.05; **p < 0.01. All data were represented as mean ± SEM. Unpaired *t*-test for C, E, F.

### GE enhances inhibitory inputs to PVT neurons

To examine GE’s effects on PVT neurons, we performed the whole-cell current-clamp recording of PVT neurons (*Figure 6A*) and bath application of GE (*Figure 6B*). The 0.3 mM GE showed a tendency to hyperpolarize the membrane potential of PVT neurons (*Figure 6C*), and 1 mM GE markedly hyperpolarized the membrane potential (*Figure 6D*). Furthermore, 1 mM GE dramatically decreased the numbers of action potentials in depolarizing step-current injections compared with the corresponding baseline (*Figure 6E*). To better illustrate GE’s effects, we displayed the activity of a representative neuron before and after 1 mM GE (*Figure 6F-I*). The neuron showed a typical action potential threshold in 50 pA depolarizing ramp-current and stable firing in 100 pA depolarizing step-current (*Figure 6F and H*). The membrane potential was significantly hyperpolarized (from −43 mV to −63 mV) and failed to generate an action potential in 200 pA depolarizing ramp-current and 100 pA depolarizing step-current pulses after GE application (*Figure 6G and I*). These results indicate that GE hyperpolarizes PVT neurons.

**Figure 6.**
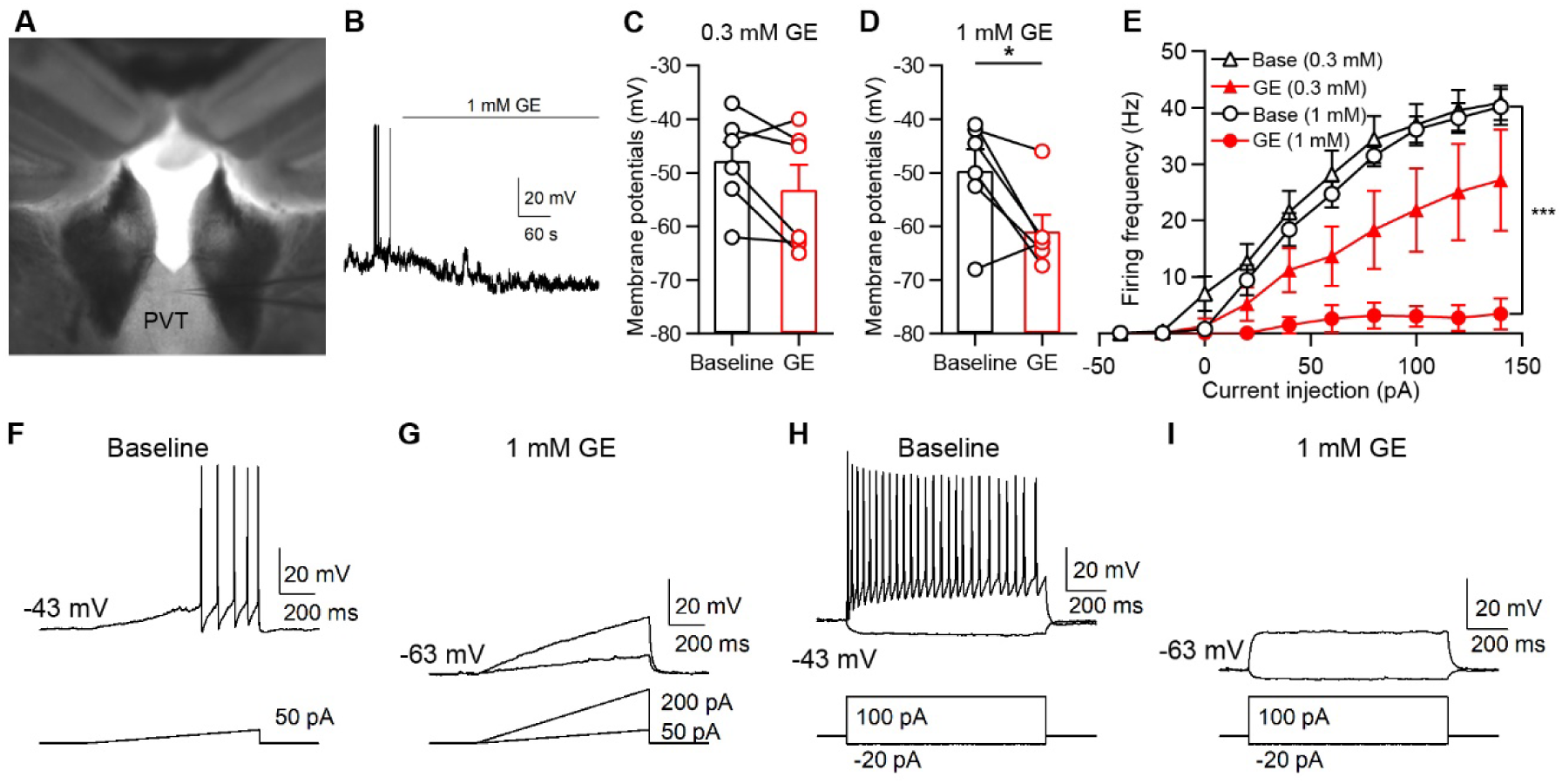
GE suppresses the activity of PVT neurons. (A) Bright field inverted microscope image of recording in PVT. (B) Whole-cell current-clamp recording (IC = 0 pA) of a representative PVT neuron with bath application of 1 mM GE. (C) The membrane potentials were hyperpolarized after application of 0.3 mM GE, *n* = 6 neurons. (D) The membrane potentials were hyperpolarized after the application of 1 mM GE, *n* = 6 neurons. (E) Summary of data showing the effect of GE (0.3 mM and 1 mM) on step-current injection-evoked spike firings of PVT neurons, *n* = 6 neurons. (F and G) A representative neuron showed GE suppressed the spike firing in response to a depolarizing ramp-current injection. (H and I) The same representative neuron showed GE suppressed the spike firing in response to a depolarizing step-current injection. *p < 0.05; ***p < 0.001. All data were represented as mean ± SEM. Paired *t*-test for D. Two-way ANOVA with Bonferroni post hoc test for E.

We further examined the detailed mechanisms of GE’s effect on PVT neurons. The suppression of PVT neurons could be due to the suppression of excitatory inputs or the enhancement of inhibitory inputs. To test these possibilities, we performed whole-cell voltage-clamp recording for PVT neurons. First, we voltage-clamped the membrane potentials of PVT neurons at −70 mV and observed the effect of 1 mM GE on spontaneous excitatory postsynaptic currents (sEPSCs) in PVT neurons. Bath application of 1 mM GE slightly increased the frequency of sEPSCs (*Figure 7–figure supplement 1A*), but not the amplitude or half-width (*Figure 7–figure supplement 1B-C*). We also analyzed the holding currents at −70 mV and found that GE did not affect the holding currents (*Figure 7–figure supplement 1D*). These results showed that GE did not affect the excitatory inputs of PVT neurons.

Next, we voltage-clamped the membrane potentials of PVT neurons at 0 mV and examined the GE’s effect on spontaneous inhibitory postsynaptic currents (sIPSCs) in PVT neurons (*Figure 7A*). Bath application of 1 mM GE did not affect the sIPSCs frequency and amplitude (*Figure 7B-C*) but increased the half-widths (*Figure 7D*). Similar results were observed in the 0.1 mM GE and 0.3 mM GE group (*Figure 7–figure supplement 1E-J*), indicating sIPSCs duration was prolonged after GE application. It is worth noting that GE application increased the holding currents at 0 mV dose-dependently (*Figure 7E*), indicating that GE induces tonic currents. According to the Cl^−^ concentrations in the internal solution (4 mM) and the ACSF solution (136.5 mM), these current analysis results suggest that GE might induce tonic Cl^−^ influx in PVT neurons.

**Figure 7.**
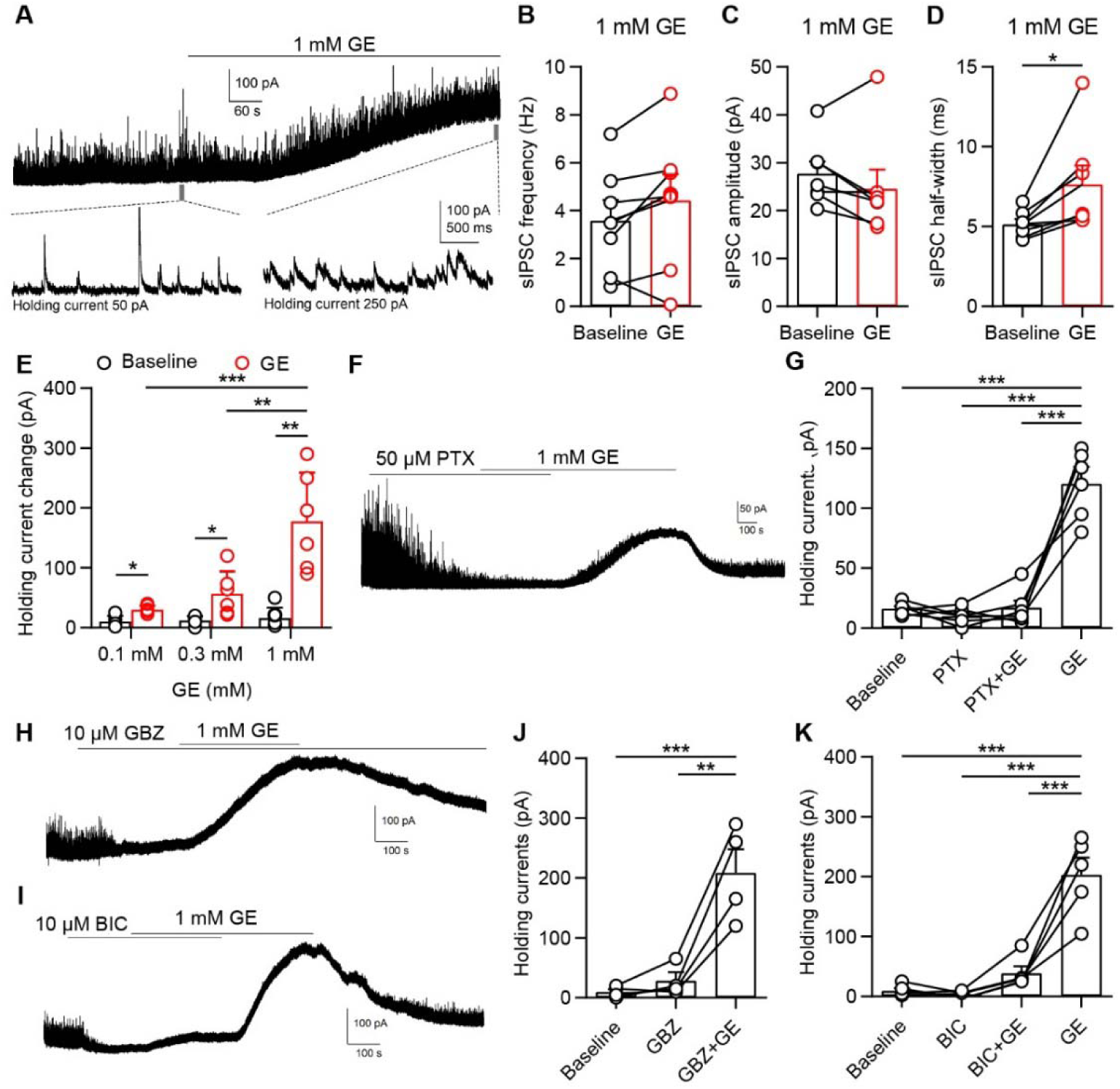
GE enhances inhibitory inputs to PVT neurons. (A) Whole-cell voltage-clamp recording (VC = 0 mV) of a representative PVT neuron. Top, bath application of 1 mM GE. Bottom, two 3-seconds time windows enlarged from before and after application of 1 mM GE (indicated by gray bars in top panel). (B-D) Frequency (B), amplitude (C), and half-width (D) of sIPSCs before and after 1 mM GE, *n* = 7 neurons. (E) Holding currents at 0 mV before and after 0.1 mM GE, 0.3 mM GE and 1 mM GE, *n* = 6 neurons. (F) A representative neuron showed picrotoxin (50 μM) blocked GE-induced tonic current in PVT neurons (VC = 0 mV). (G) Statistical comparison of holding currents changes, *n* = 6 neurons. (H) A representative neuron showed gabazine (GBZ, 10 μM) did not block GE-induced tonic current in PVT neurons (VC = 0 mV). (I) A representative neuron showed bicuculline (BIC, 10 μM) blocked GE-induced tonic current in PVT neurons (VC = 0 mV). (J) Statistical comparison of holding currents changes in GBZ experiments, *n* = 4 neurons. (K) Statistical comparison of holding currents changes in BIC experiments, *n* = 5 neurons. *p < 0.05, **p < 0.01, ***p < 0.001. All data were represented as mean ± SEM. Paired t-test for D. Two-way ANOVA with Bonferroni post hoc test for E. One-way ANOVA with Bonferroni post hoc test for G, J, K. The following figure supplement is available for figure 7: **Figure supplement 1.** The effect of GE on sEPSCs, sIPSCs, and holding currents in PVT neurons. **Figure supplement 2.** Molecular Docking of GE to GABA_A_Rβ3.

### GE induces tonic inhibition via GABA_A_ receptors

GABA_A_ receptors are critical targets for various sedatives and anesthetics (Kim et al., 2020). A computational study revealed that the GABA_A_ receptor α1 and β1 subunits might also be putative targets of GE (Zhang et al., 2019). We speculated that GE might induce tonic currents by acting on GABA_A_ receptors. Picrotoxin (PTX) is an open channel blocker of GABA_A_ receptors and blocks synaptic GABA_A_ currents as well as tonic extrasynaptic currents (Wlodarczyk et al., 2013). We applied picrotoxin (50 μM) before 1 mM GE and found that GE failed to induce tonic currents in the presence of picrotoxin. We further washed out picrotoxin and then observed the GE-induced tonic currents (*Figure 7F*). The statistical data showed that picrotoxin completely blocked the GE-induced holding current change (*Figure 7G*).

Gabazine (GBZ, also known as SR-95531) and bicuculline (BIC) are competitive GABAA receptor antagonists. Previous studies have found that gabazine did not prevent tonic inhibition, while bicuculline blocked tonic currents in hippocampal neurons and spinal dorsal horn neurons (Bai et al., 2001; Maeda et al., 2010; Wlodarczyk et al., 2013). We used gabazine and bicuculline to further investigate the potential mechanism of GE acting on GABA_A_ receptors. First, we recorded PVT neurons and perfused them in 10 μM gabazine. We found that the sIPSCs were blocked while the holding current was unaffected. The application of GE still induced tonic currents remarkably in the presence of gabazine (*Figure 7H and J*). When 10 μM bicuculline was applied before GE, the GE-induced tonic currents were dramatically blocked, while washing out of bicuculline led to tonic currents (*Figure 7I and K*). These data indicate that GE-induced tonic inhibition is blocked by bicuculline but not gabazine. Taken together, these results demonstrated that GE induces tonic inhibition by acting on GABA_A_ receptors.

It is reported that the binding of bicuculline further allosterically closed two β-α and one α-β interface as well as conferring rigid-body subunit transformation (Kim et al., 2020). We further used molecular docking and molecular simulations to study the interaction between GE and GABA_A_R subunits α1-6 and β1-3, and found that GABA_A_Rβ3 has the best binding energy (*Figure 7–figure supplement 2A*). The optimal crystal structure of mouse GABA_A_Rβ3 fulfilled the probability density function energy requirements through the Ramachandran plot test (*Figure 7–figure supplement 2B*). The binding sites of GE to GABA_A_Rβ3 were revealed in *Figure 7–figure supplement 2C*. This precise match suggested that GE might interact with the GABA_A_Rβ3 activated pocket (Leu435). The molecular docking result informed that GE was a potential ligand for interacting with GABA_A_Rβ3.

## Discussion

In this study, we found that GE had locomotor sedating, anti-convulsant, and hypnotic effects. Next, we observed that the systemic administration of GE suppressed the activity of the PVT. Microinjection of GE in the PVT could facilitate propofol-induced anesthesia and suppress locomotion. Furthermore, we found that GE induced dramatic membrane hyperpolarization and suppressed PVT neurons, mainly by prolonging the spontaneous inhibitory postsynaptic currents and inducing tonic inhibitory currents via GABA_A_ receptors. In conclusion, we showed that GE plays a vital role in inducing sedation in mice. These findings provide a potential candidate for the development of sedatives and anesthetics.

### Dissection of GE’s sedative effects by multiple behavioral tests

Geraniol is abundant in the essential oils extracted from lemongrass, rose, lavender, and other aromatic plants, and it is widely used in the fragrance market due to its rose-like odor (Chen and Viljoen, 2010; Zhang et al., 2019). Recent studies found that GE treatment relieved pathological pain in mice (La Rocca et al., 2017; Lv et al., 2017). Additionally, several studies have shown that GE could modulate sedation in rodents or aquatic species (Can et al., 2019; Medeiros et al., 2018). They found GE could increase barbiturate-induced sleeping time in rats and conjectured that these effects were related to GE’s depressant effect on the central nervous system (Medeiros et al., 2018). In our study, we adopted multiple behavioral tests to dissect the role of GE in sedation.

Inhaled therapies are the cornerstone of treatment in clinics, and aromatherapy has great market potentials. Inhalation is an ideal administration route for essential oils according to the bioavailability, toxicity, and therapeutic efficacy. Previous studies have used inhalation of a single component or complex of essential oils to study the sedative effects or anti-inflammatory effects in mice (Linck et al., 2009; Ueno-Iio et al., 2014). Volatile anesthetics, such as sevoflurane and isoflurane, are ideal for direct vaporization. For sevoflurane, the boiling point is 58.5 °C, and vapor pressure is 193 mm Hg at 25 °C. The boiling point is 48.5 °C, and vapor pressure is 330 mm Hg at 25 °C for isoflurane. GE is a low molecular weight monoterpenoid with liposolubility. However, the boiling point for GE is 227.5 °C, and vapor pressure is 0.03 mm Hg at 25 °C. So direct vaporization of GE by airflow is far less efficient than sevoflurane or isoflurane. In our experiments, we used ultrasonic nebulization equipment because it could control the GE concentration (1.5% in ddH_2_O) and nebulization rate (65 ml per 40 minutes). Inhalation of GE suppressed the locomotion and facilitated propofol-induced anesthesia, indicating its locomotor sedating effect and hypnotic effect. Systemic administration of GE by intraperitoneal injection showed similar effects. These data were consistent with the previous study in rats (Medeiros et al., 2018). Since GE is recognized as a safe flavor ingredient by FDA, GE has both clinical and economic value in the future.

Another characteristic of sedatives is the anti-convulsant effect. Previous studies have shown that dehydrofukinone, isolated from *Nectandra grandiflora (Lauraceae)* essential oil, could delay the onset of generalized tonic-clonic seizures but did not alter the severity of seizures (Garlet et al., 2017). Here we found that GE significantly decreased the severity of seizures in the 3-5 minutes, which is the most severe period during the 15 minutes recording. These data indicate that GE has a robust anti-convulsant effect in mice. Moreover, we observed that GE groups (especially in the dose of 100 mg/kg) showed a slightly increased severity score after 5 minutes, which might be caused by the metabolized reduction of GE and the rebound of seizures. In the 200 mg/kg GE group, the rebound of seizures was not significant.

We also observed the hypnotic effect of GE by recording the LORR/RORR in different anesthetics models, including intravenous injection of propofol, intraperitoneal injection of pentobarbital sodium, and inhalation of isoflurane. Importantly, we found that the RORR in the 200 mg/kg GE + 15 mg/kg propofol group was comparable to the 20 mg/kg propofol group, indicating that GE could reduce the dosage of propofol. Multimodal general anesthesia is a modern management strategy. It is postulated that the use of more agents at smaller doses could maximize desired effects while minimizing side effects (Brown et al., 2018). GE’s hypnotic effect might be taken into consideration for choosing the drug combinations in the future.

We further evaluated the pattern of brain waves in GE-treated mice. The PSD of delta waves (0.5-4 Hz) and theta waves (4-7 Hz) were enhanced after GE injection, in accordance with the previous studies that GE could increase delta wave power in rats (Medeiros et al., 2018). Generally, the EEG is dominated by the delta and theta waves during the non-rapid eye movement (NREM) sleep (Scammell et al., 2017). Other studies showed increased delta waves and theta waves in multiple cortexes during the slow-wave sleep in rats (Jing et al., 2016). The EEG alterations indicate that GE has a hypnotic effect consistent with the LORR/RORR behavioral results.

### GE suppressed the PVT activity by enhancing inhibitory inputs via GABA_A_ receptors

A recent study reported that the PVT is a critical node for controlling wakefulness in rodents (Ren et al., 2018). They found that activation of the PVT enhanced wakefulness, and suppression of the PVT reduced wakefulness. Here we observed that GE suppressed the activity of PVT from c-Fos staining results. Moreover, *in vivo* fiber photometry results showed that the calcium signal of the PVT was significantly decreased after GE. These data confirmed that GE could suppress the activity of PVT. Further microinjection of GE into the PVT facilitated propofol-induced anesthesia and reduced locomotion, indicating that the PVT is a crucial brain region responsible for GE’s sedative effects.

The following whole-cell recording results showed that GE remarkably suppressed the activity of PVT neurons. The frequency of sEPSCs was slightly increased, while the amplitude or the half-width of sEPSCs was not affected. The increase in sEPSCs frequency might be caused by GE-induced disinhibition of presynaptic excitatory neurons. We speculated that further experiments with sodium channel blocker tetrodotoxin (TTX) should block action potentials and eliminate the disynaptic effect. Meanwhile, the frequency and amplitude of sIPSCs were unaffected. Only the half-width was consistently increased, indicating a change in the kinetics of channel closing. This phenomenon is similar to the propofol-induced reduction in the decay rate of sIPSCs (Drexler et al., 2016; Orser et al., 1994). Recent studies have revealed that the PVT receives GABAergic inputs from many brain nuclei, including the reticular thalamic nucleus, zona incerta, and hypothalamic arcuate nucleus (Betley et al., 2013; Lee et al., 2019; Zhang and Van Den Pol, 2017). Whether GE facilitates inhibitory inputs from specific nuclei or enhances the overall GABAergic tone is still unknown.

Sedatives, such as propofol and midazolam, can induce tonic currents in hippocampal neurons or somatosensory cortex neurons (Bai et al., 2001; Yamada et al., 2007). The tonic inhibition has a profound influence on neural excitability, synaptic plasticity, neurogenesis, and network oscillations (Duveau et al., 2011; Ge et al., 2006; Martin et al., 2010; Pavlov et al., 2009). It is mainly induced by the activation of GABA_A_ receptors distributed extrasynaptically, where they are exposed to fluctuating but low concentrations of GABA (Franks, 2008). Several studies have shown that tonic inhibition is mediated by constitutively active GABA_A_ receptors in the absence of GABA (O’Neill and Sylantyev, 2018; Pavlov et al., 2009).

A previous study showed that bicuculline and gabazine acted as competitive GABA_A_ receptor antagonists, but only bicuculline blocked tonic currents (Bai et al., 2001). Similarly, in our results, bicuculline blocked GE-induced sIPSCs and also tonic inhibition, while gabazine only blocked sIPSCs. A recent study revealed that cryo-electron microscopy structures of the α1β2γ2 GABA_A_ receptor bound to general anesthetics (Kim et al., 2020). The binding of bicuculline allosterically closes three of the anesthetic pockets, including two β-α interfaces and one α-β interface, as well as inducing rigid-body subunit transformations (Kim et al., 2020; Ueno et al., 1997). Another work using the GABA_A_ receptor reported that the binding of bicuculline at the orthosteric sites prevents closure of the β3-α1 interfaces and rotation of the extracellular domains, and also stabilizes transmembrane domains in the closed state (Masiulis et al., 2019). We carried out molecular docking and molecular simulations to study the interaction between GE and GABAAR subunits α1-6 and β1-3, and found that GABA_A_Rβ3 has the best binding energy and Leu435 might be an interacted pocked. We conjecture that GE might bind to the β3 subunit (other binding sites are not excluded), inducing allosteric activation of GABA_A_ receptors and tonic inhibition. These binding sites or the transformation change could be blocked by bicuculline but not gabazine.

The metabotropic GABA_B_ receptor is a G protein-coupled receptor that mediates slow and prolonged inhibitory neurotransmission in the brain. The GABA_B_ receptors require two distinct subunits (GABA_B_1 and GABA_B_2) and locate both pre-synaptically and post-synaptically. They are coupled to K^+^ and Ca^2+^ channels, and the activation of GABA_B_ receptors leads to a variety of effects, such as inhibition of transmitter release and neuronal hyperpolarization (Bowery, 2006; Chalifoux and Carter, 2011). The ligand-binding mechanism and conformational change of GABA_B_ receptors are different from those of GABA_A_ receptors (Frangaj and Fan, 2018), and we cannot exclude the possible activation of GABA_B_ receptors by GE. Furthermore, GE might also affect calcium and potassium channels, including voltage-dependent potassium ion channels (de Menezes-Filho et al., 2014; Ye et al., 2019). Further structure and mutation studies are needed to reveal the detailed mechanisms of these processes.

Consequently, our findings identify that GE enhances inhibitory inputs to PVT neurons, induces sedation in mice, and suggest a potential candidate for further development of sedatives and anesthetics. An attempt to use GE or combination GE with anesthetics in the clinic could be very promising.

## Materials and methods

### Animals

Postnatal days (P) 30 and 60 male *C57BL/6J* mice purchased from SLAC laboratory (Shanghai) were used for experiments. For slice recording experiments, P30 mice were used. All mice were raised on a 12-hour light/dark cycle (lights on at 7:00 am) with *ad libitum* food and water. All behavioral tests were carried out during the light phase. All animal experiment procedures were approved by the Animal Care and Use Committee of Shanghai General Hospital (2019AW008) and Animal Care and Use Committee of Fudan University School of Basic Medical Sciences (20180511-001).

### Systemic administration of GE

For nebulization of GE (Sigma, 163333), 1.5 ml GE was mixed with 100 ml ddH_2_O (vortex thoroughly). A nebulizer (yuwell, 402B) was used to control the nebulization rate (65 ml per 40 minutes). Mice were kept in a transparent plexiglass induction chamber (30 x 16 x 20 cm), and GE (1.5% in ddH_2_O, 65 ml) was nebulized for 40 minutes. Control mice were kept in the identical chamber with nebulized water (65 ml) for 40 minutes.

For intraperitoneal injection (i.p.), GE was diluted in corn oil (C805618, MACKLIN) by vortex thoroughly. Mice were intraperitoneally injected with GE (100, 200, or 400 mg/kg in weight, i.p.). For the control group, mice were intraperitoneally injected with the same volume of corn oil.

### Implantation and microinjection in PVT

Mice were anesthetized by vaporized isoflurane (induction, 2%; maintenance, 1.5%) and head-fixed in a mouse stereotaxic apparatus (RWD Life Science Co.). A cannula (outside diameter 0.41 mm, internal diameter 0.25 mm; length 6 mm, RWD Life Science Co.) was implanted into the PVT (AP −1.46 mm, ML 0 mm, DV −2.90 mm). Two weeks later, 200 nl of GE (1 mM in ACSF) or ACSF was microinjected into the PVT at a rate of 200 nl/min for 1 minute by standard infuse/withdraw pump (Harvard Apparatus). The injection needle was stayed for an additional 2 minutes to allow drug diffusion.

### Open field test

Mice were tested their locomotor activity in plexiglass enclosures (40 x 40 x 40 cm). Fifteen minutes after GE injection (i.p., or microinjection to PVT), mice were placed in the center of the box and were videotaped individually for 10 minutes. The center area was defined as centric 20 x 20 cm. The track was analyzed by AniLab software (Ningbo AnLai). Total distance, move duration, and velocity were analyzed.

### PTZ-induced Seizure

PTZ (P6500, Sigma) was dissolved in saline at a concentration of 6 mg/ml. For the induction of seizures, PTZ was administered intraperitoneally at 55 mg/kg body weight. Animals were monitored for 15 minutes after the injection. Behavioral responses were recorded using a video camera and scored at every 1 minute as follows: no abnormal behavior (0), reduced motility and prostate position (1), partial clonus (2), generalized clonus including extremities (3), tonic-clonic seizure with rigid paw extension (4) and death (5) (Li et al., 2012; Takahashi et al., 2012).

### Propofol-induced general anesthesia

Fifteen minutes after GE injection (i.p., or microinjection into PVT), the mice were anesthetized by propofol (20 mg/kg or 15 mg/kg, i.v.). The interval from propofol injection to loss of righting reflex (LORR) was measured as LORR time. The LORR of propofol-induced anesthesia is less than 5 seconds in our experiments. The interval from loss of right reflex to return of righting reflex (RORR) was measured as RORR time. Propofol was purchased from Beijing Fresenius Kabi Co., Ltd.

### Pentobarbital sodium-induced general anesthesia

Fifteen minutes after GE injection, the mice were anesthetized by pentobarbital sodium (50 mg/kg, i.p., Merck). And the LORR time and RORR time were measured.

### Isoflurane-induced general anesthesia

Isoflurane (RWD Life Science Co.) was delivered in 400 ml/min using an isoflurane vaporizer (MSS, UK) and an open-circuit rodent anaesthesia system. Fifteen minutes after GE injection, mice were placed into induction chambers prefilled with 2 vol% isoflurane. The interval between when anesthetic administration was initiated and LORR time was then recorded. After LORR, isoflurane 1.5 vol% was continued for 10 minutes to ensure equilibration. Then, isoflurane administration was withdrawn, and the mice were considered to have recovered the righting reflex if they could turn themselves to the prone position. The interval between discontinuation of anesthetic and the return of righting reflex was determined as the time of RORR.

### Morris water maze (MWM)

MWM was carried out to assess spatial learning and memory function (Vorhees and Williams, 2006). The water maze was 120 cm cylindrical tank in diameter with a 10 cm platform in diameter, filled with opaque water obscuring the platform (water is 2 cm above the platform height). The platform was located in the center of one quadrant. Four visual cues were posted around the tank wall. An overhead camera and AniLab software (AniLab Tech, Ningbo, China) were used to track and analyzed the movement of animals, including latency, swimming distance, crossing number, and speed.

Twenty-four hours after corn oil (i.p.), propofol (20 mg/kg, i.v.), or propofol (20 mg/kg, i.v.) with GE (200 mg/kg, i.p.) were injected, mice were trained for 5 consecutive days (Day 1-5). During each acquisition day, mice had four trials (start from different quadrants, 30 minutes intervals). Mice were given 60 seconds to find the platform and allowed to stay for 15 seconds. If a mouse could not find the platform within 60 seconds, it was gently guided to the platform and allowed to remain there for 15 seconds. On the probe trial on Day 6, mice had a 60 seconds test for traveling in the tank without a platform.

### Fear conditioning test

Fear conditioning was conducted in a conditioning chamber (25 x 30 x 20 cm, ENV-008, Med Associates). The chamber was located in a sound-attenuating box (NIR-022MD, Med Associates). Electric footshock unconditioned stimulus (US) was produced by a shock generator (ENV-414S, Med Associates). Auditory conditioned stimulus (CS) was produced by a speaker (ENV-224AM, Med Associates).

Twenty-four hours after the drugs were injected, mice were performed the fear conditioning test for three consecutive days (Day 1-3) (Shoji et al., 2014). On Day 1 (habituation), the mice were habituated to the fear condition chambers. After 2 minutes of exploration, three tones (75 dB, 4k Hz, 30 seconds duration) separated by a variable interval with a range of 60-120 seconds and an average of 90 seconds were delivered. On Day 2 (conditioning), mice received five trials of the CS paired with the US separated by 90 seconds. The CS was 75 dB, 4 kHz, 30 seconds duration, co-terminated with a footshock US (0.6 mA, 1-second duration). After the last CS/US conditioning, the mice were kept in the conditioning chamber for another 60 seconds before being returned to the home cages. On Day 3, the mice were placed into a modified chamber to perform the tone cue test. The chamber was modified by replacing its metal floor with a plastic floor, adding a black triangular ceiling. Mice were placed in the altered chamber for 3 minutes to measure the freezing level in the altered context. Then, a tone (75 dB, 4 kHz) was delivered for 2 minutes.

The behavior of the mice was recorded and analyzed with the Video Freeze software (Med Associates, St Albans, VT). Motionless bouts lasting more than 0.5 seconds were defined as freeze. On Day 1, the freezing percentages in the first 2 minutes exploration period and during the three tones were defined as the baseline freezing percentages to the environment and the tone. On Day 2, the freezing percentages in the first 29 seconds period during each tone (except the shock duration) were summarized as an indication of fear memory acquisition. On Day 3, the freezing percentages were calculated in the 2 minutes cue test.

### Electroencephalogram (EEG)

Mice were anesthetized by vaporized isoflurane (induction, 2%; maintenance, 1.5%) and head-fixed in a mouse stereotaxic apparatus. Erythromycin ointment was applied to the eyes of mice to avoid corneal drying. The scalp was shaved, and the skull was exposed under antiseptic conditions. Four copper screws were installed at the 1 mm anterior to bregma and 1 mm anterior to the lambda with 1.5 mm lateral on both sides of the skull without penetrating the underlying dura. Then the insulated wires from the EEG, reference, and ground electrodes were welded to the screws. The electrode apparatus was fixed on the skull using 3M tissue glue followed by dental cement.

The mice were placed in the behavior chamber for at least 30 minutes a day for 3 consecutive days before the experiment to ensure that the mice acclimate to the environment. The video camera was used to consecutively record the behaviors of the mice. The headstage and EEG electrodes were gently connected. The mice were adapted for 30 minutes and followed by a 10-minute baseline recording. Cortical EMG signal was recorded using a Zeus system (Zeus, Bio-Signal Technologies: McKinney, TX, USA), and LFP signals were filtered online at 200 Hz (3 kHz sampling rate). Then the mice were injected intraperitoneally with corn oil or geraniol, and the EEG signal was recorded lasting for 30 minutes.

Welch’s averaged periodogram method with a 1024 ms nonoverlapping Hanning window (Nonuniform fast Fourier transform, NFFT = 2048) was used to perform a power spectral density analysis on local field potential (LFP) signals. A time-frequency diagram of LFP was performed using a short-time Fourier transform in overlapping 512 ms Hanning window with a step size of 50 ms (NFFT = 512).

### Surface electrocardiography (ECG)

Once the righting reflex was lost after propofol injection, the mice were placed in the supine position on a rodent surgical monitor (Indus Instruments). The limbs of the mouse were closely attached to the electrode plate heating pad with conductive glue. Respiratory rate and peripheral oxygen saturation (SpO_2_) were measured for 2-3 minutes. Data between 60 sec to 120 sec after LORR were extracted every 10 sec by offline analysis.

### Histology

Animals were deeply anesthetized with vaporized sevoflurane and transcardially perfused with 20 ml saline, followed by 20 ml paraformaldehyde (PFA, 4% in PBS). Brains were extracted and soaked in 4% PFA at 4°C for a minimum of 4 hours and subsequently cryoprotected by transferring to a 30% sucrose solution (4°C, dissolved in PBS) until brains were saturated (for 36-48 hours). Coronal brain sections (40 μm) were cut using a freezing microtome (CM1950, Leica). The slices were collected and stored in PBS at 4°C until immunohistochemical processing.

The brain sections undergoing immunohistochemical staining were washed in PBS for 3 times (10 minutes each time) and incubated in a blocking solution containing 0.3% TritonX-100 and 5% normal donkey serum (Jackson ImmunoResearch, USA) in PBS for 1 hour at 37°C. Sections were then incubated (4°C, 24 hours) with primary antibodies anti-rabbit c-Fos (1:4000, ab190289, Abcam) dissolved in 1% normal donkey serum solution. Afterward, sections were washed in PBS for 4 times (15 minutes each time), then incubated with secondary antibodies Alexa Flour 488 conjugated donkey anti-rabbit IgG (1:800, Jackson) for 2 hours at room temperature. Nuclei were stained with DAPI (Beyotime, 1:10000) and washed three times with PBS. The photofluorograms were taken by the Leica DMi8 microscope. The photomicrographs were further processed by Fiji.

### Fiber photometry

*In vivo* fiber photometry experiments were performed as previously described *(Zhu et al., 2020)*. The AAV2/8-hSyn-GCaMP6s virus (200 nl, 4 x 10^12^ v.g./ml, S0225-8, Taitool Bioscience) was injected into the PVT nucleus (antero-posterior, AP, -1.34 mm, medio-lateral, ML, 0 mm, dorsal-ventral, DV, -2.9 mm) of the WT mice, the optic fiber was implanted above the PVT. After three weeks for virus expression, the mice were gently handled to be familiar with the calcium signal recording system (Thinker-Biotech). The LED intensity was 10-15 μW, and the fluorescence signal was recorded at 50 Hz. We defined a 2-minute time window before GE or corn oil injection as the baseline period. And we defined a 2-minute time window 10 minutes after GE or corn oil injection as the post-injection period.

### Brain slice electrophysiology

P30 Mice were anesthetized with isoflurane and perfused transcardially with an ice-cold cutting solution containing (in mM): sucrose 213, KCl 2.5, NaH_2_PO_4_ 1.25, MgSO_4_ 10, CaCl_2_ 0.5, NaHCO_3_ 26, glucose 11 (300-305 mOsm). Then the brain was rapidly dissected, and coronal slices (280 μm) were sectioned. Slices were transferred into holding chamber and incubated in 34°C artificial cerebrospinal fluid (ACSF) containing (in mM): NaCl 126, KCl 2.5, NaH_2_PO_4_ 1.25, MgCl_2_ 2, CaCl_2_ 2, NaHCO_3_ 26, glucose 10 (300-305 mOsm). After 30 minutes of recovery, the holding chamber with the slices was transferred to room temperature (22-24°C). Both cutting solution and ACSF were continuously bubbled with 95% O_2_/5% CO_2_. Then, slices were placed on glass coverslips coated with poly-L-lysine and submerged in a recording chamber. All experiments were performed at near-physiological temperatures (30-32°C) using an in-line heater (Warner Instruments) while perfusing the recording chamber with ACSF at 3 ml/min using a pump (HL-1, Shanghai Huxi). Whole-cell patch-clamp recordings were made from the target neurons under IR-DIC visualization and a CCD camera (Retiga ELECTRO, QIMAGING) using a fluorescent Olympus BX51WI microscope. Recording pipettes (2-5 MΩ; Borosilicate Glass BF 150-86-10; Sutter Instrument) were prepared by a micropipette puller (model P97; Sutter Instrument) and backfilled with potassium-based internal solution containing (in mM) K-gluconate 130, MgCl_2_ 1, CaCl_2_ 1, KCl 1, HEPES 10, EGTA 11, Mg-ATP 2, Na-GTP 0.3 (pH 7.3, 290 mOsm) or cesium-based internal solution contained (in mM) CsMeSO_3_ 130, MgCl_2_ 1, CaCl_2_ 1, HEPES 10, QX-314 2, EGTA 11, Mg-ATP 2, Na-GTP 0.3 (pH 7.3, 295 mOsm). Biocytin (0.2%) was included in the internal solution.

In dissecting the effect of GE on the firing rate activity of PVT neurons, neurons in PVT were recorded in I = 0 mode (potassium-based internal solution). After stable recording spontaneous firing for 2 to 3 minutes, 1 mM GE was bath applied for 8 to 10 minutes. One-minute baseline and one-minute post-GE (8 or 9 minutes after application of GE) were analyzed.

In dissecting the effect of GE on spontaneous excitatory currents (sEPSCs) or spontaneous inhibitory currents (sIPSCs) of PVT neurons, neurons in PVT were recorded in voltage-clamp mode (cesium-based internal solution, holding voltage -70 mV or 0 mV). After stable recording for 2 to 3 minutes, 0.1 mM, 0.3 mM, or 1 mM GE was bath applied for 8 to 10 minutes. One-minute baseline and one-minute post-GE (8 or 9 minutes after application of GE) were analyzed.

In dissecting the effect of picrotoxin, gabazine, or bicuculline on GE-induced holding current at 0 mV, neurons in PVT were recorded in voltage-clamp mode (cesium-based internal solution, voltage clamp at 0 mV). After stable recording for 2 to 3 minutes, 50 μM picrotoxin, 10 μM gabazine, or 10 μM bicuculline was bath applied for 5 to 8 minutes. Then 1 mM GE was bath applied together with different GABA receptor antagonists for 8 to 10 minutes. Different wash-out sequences were used according to the effect of GE on PVT neurons in the presence of different GABA receptor antagonists. For picrotoxin and bicuculline experiments, picrotoxin was first washed out, then GE was washed out. For the gabazine experiment, GE was first washed out, and then gabazine was washed out.

Picrotoxin was purchased from Tocris Bioscience (1128). Gabazine was purchased from MedChemExpress (HY-103533). Bicuculline methiodide was purchased from Abcam (ab120108). All other chemicals were obtained from Sigma.

### Electrophysiology data acquisition and analysis

Data Acquisition and analysis were performed as described previously (Zhou et al., 2017). Voltage-clamp and current-clamp recordings were carried out using a computer-controlled amplifier (MultiClamp 700B; Molecular Devices). During recordings, traces were low-pass filtered at 4 kHz and digitized at 10 kHz (DigiData 1550B1, Molecular Devices). Data were acquired by Clampex 10.6. Data were filtered using a low-pass-Gaussian algorithm (-3 dB cutoff frequency = 1000 Hz) in Clampfit 10.6 (Molecular Devices). Extremely small, low signal-to-noise ratio and unreliable responses were regarded as no response. The frequency, amplitude, half-width of sEPSCs or sIPSCs were analyzed by event detection using self-defined templates in Clampfit 10.6.

### Homology modeling and molecular docking

Homology modeling and molecular docking were performed as described previously (Wang et al., 2018). On account of the crystal structure of mouse GABA_A_R subunits that were not analyzed in RCSB Protein Data Bank (PDB), we downloaded the amino acid sequence of each GABA_A_R subunit from the UniProtKB database (http://www.uniprot.org/) to establish the protein crystal structure. Homology modeling was applied with the SWISS-MODEL (https://www.swissmodel.expasy.org/) to obtain the structure of each GABA_A_R subunit based on suited structure. PROCHECK was used to examine the stereochemical quality of the structure obtained from SWISS-MODEL to draw the Ramachandran plot. The interactions between GE and GABA_A_R subunits were simulated via AutoDocking vina. Default AutoDocking vina parameters were applied. Among these subunits, the interaction affinity between GE and GABA_A_Rβ3 activated pocket (Leu435) was the highest. Finally, the protein-ligand complexes were viewed by Pymol.

### Quantification and statistical analysis

Software used for data analysis included: Clampfit 10.6, GraphPad Prism 6, MATLAB 2019, Fiji. Statistical detection methods include student’s unpaired *t*-test, paired *t*-test, one-way or two-way ANOVA with *post hoc* Bonferroni test. A value of P < 0.05 was considered as statistically significant, and data were shown as mean ± SEM.

## Acknowledgment

We thank Dr. Xing-Jun Liu (Shantou University Medical College, Shantou, China) for the valuable discussion. We thank Shuo Wang (Sixth People’s Hospital, Shanghai Jiao Tong University) for the technical support. This work was supported by the National Natural Science Foundation of China (No. 31900717, 81771202), the Shanghai Sailing Program (19YF1438700), Youth talent support program from China Association for Science and Technology (2019QNRC001, to D.M.), Youth talent support program from Shanghai Jiao Tong University School of Medicine (19XJ11010, to D.M.), Innovative Research Team of High-level Local University in Shanghai (to W.L. M.), Shanghai Key Laboratory of Acupuncture Mechanism and Acupoint Function (to W.L. M.). We thank Enago (http://www.enago.com/) for the English language review.

## Supplemental Information

**Figure 3–figure supplement 1.**
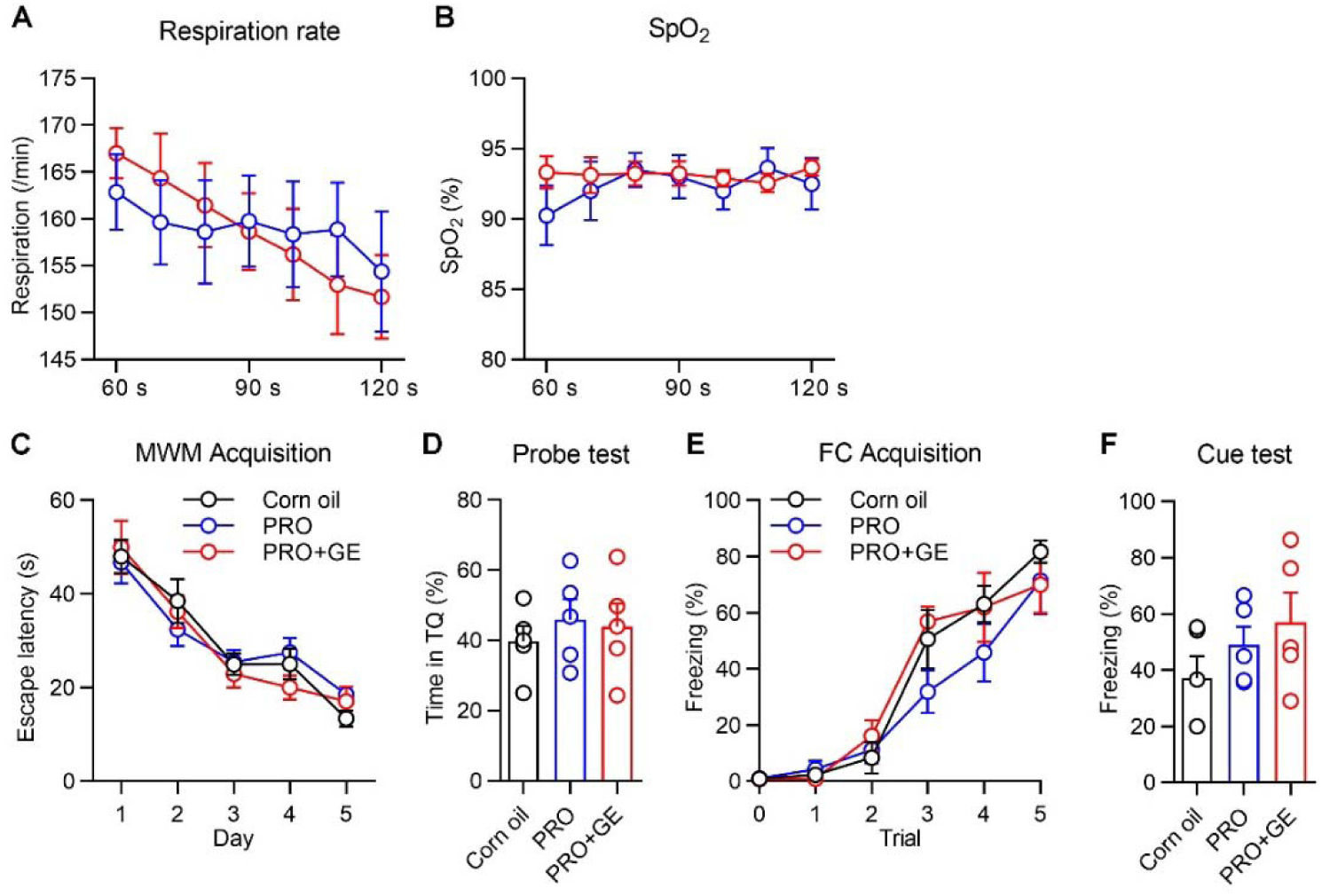
GE does not impair learning and memory in propofol-induced anesthesia of mice. (A and B) Respiratory rate (A) and oxygen saturation (B) of respiratory function before and after 1 mM GE, *n* = 8−9 mice. (C) The escape latencies during acquisition in Morris water maze test on Day 1-5, *n* = 5 mice. (D) Probe test on Day 6. (E) Percent of freezing during acquisition in fear conditioning test on Day 1, *n* = 5 mice. (F) Cue test on Day 2. All data were represented as mean ± SEM.

**Figure 7–figure supplement 1.**
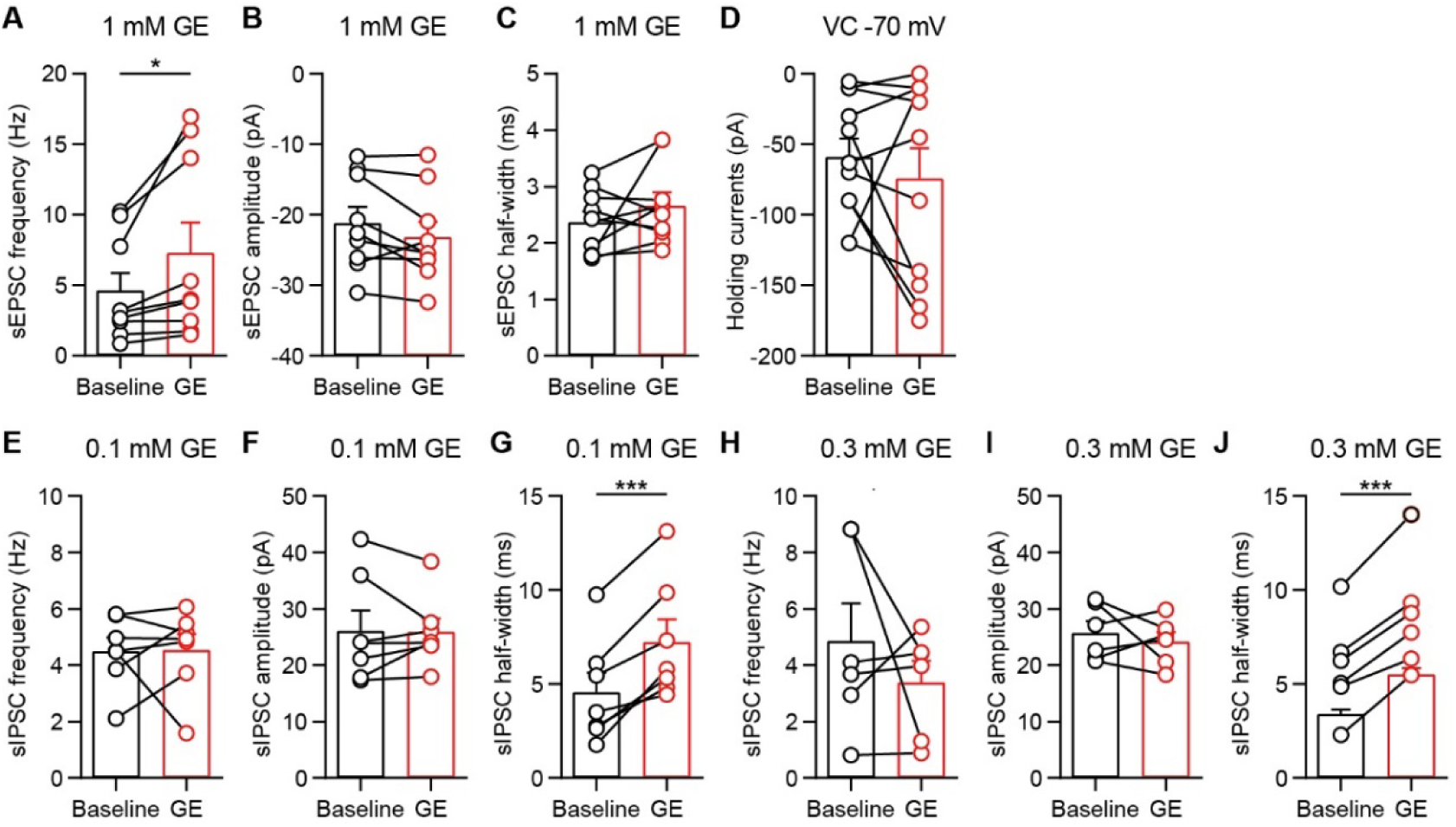
The effect of GE on sEPSCs, sIPSCs, and holding currents in PVT neurons. (A-C) Frequency (A), amplitude (B) and half-width (C) of sEPSCs before and after 1 mM GE, *n* = 9 neurons. (D) Statistical comparison of 1 mM GE-induced holding currents changes (VC = -70 mV), *n* = 11 neurons. (E-G) Frequency (E), amplitude (F) and half-width (G) of sIPSCs before and after 0.1 mM GE, *n* = 7 neurons. (H-J) Frequency (H), amplitude (I) and half-width (J) of sIPSCs before and after 0.3 mM GE, *n* = 6 neurons. *p < 0.05, ***p < 0.001. All data were represented as mean ± SEM. Paired *t*-test for A.

**Figure 7–figure supplement 2.**
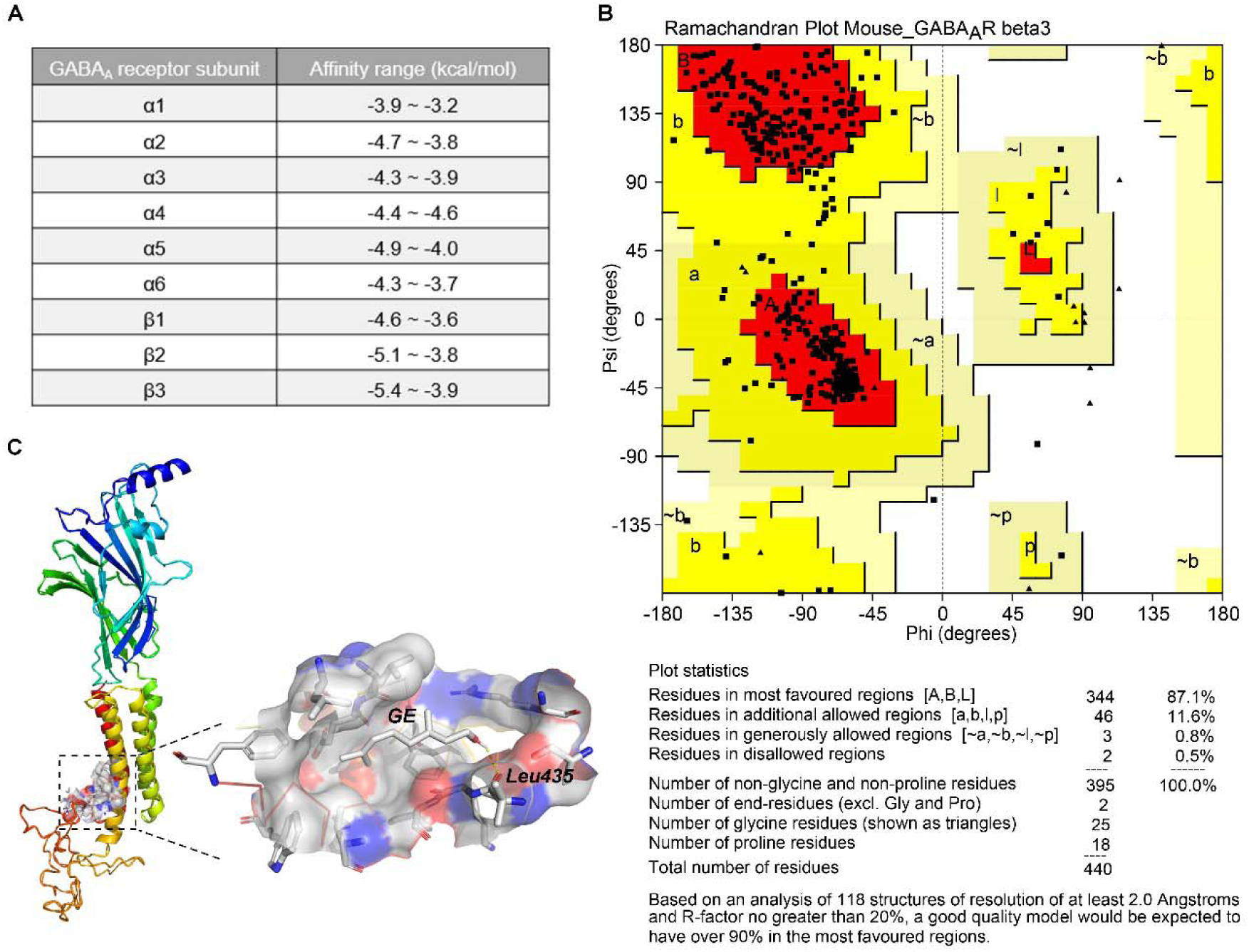
Molecular Docking of GE to GABA_A_Rβ3. (A) The top binding affinity range of GE to GABA_A_R subunits was listed. Affinity represents binding energy. (B) Ramachandran plot of mouse GABA_A_Rβ3. The different color areas of the Ramachandran plot are shaded in core (Red), allow (Yellow), generous (Light yellow), disallow (White). 87.1% of amino acid residues are located in the core area; 11.6% amino acid residues are located in the allowed area. (C) The binding sites of GE to GABA_A_Rβ3 were showed. GE interacts with residues in hydrogen bondings towards Leu435.

## Notes

### Competing Interest Statement

The authors have declared no competing interest.

